# A rationally designed inactivated *Salmonella* Typhimurium vaccine induces strong and long-lasting immune responses in pigs

**DOI:** 10.1101/2023.03.27.534319

**Authors:** Verena Lentsch, Selma Aslani, Thomas Echtermann, Swapan Preet, Elisa Cappio Barazzone, Daniel Hoces, Claudia Moresi, Dolf Kuemmerlen, Emma Slack

## Abstract

*Salmonella enterica* subspecies *enterica* serovar Typhimurium (*S*.Tm) poses a considerable threat to public health due to its zoonotic potential. Human infections are mostly foodborne, and pork and pork products are ranked among the top culprits for transmission. In addition, the high percentage of antibiotic resistance, especially in monophasic *S*.Tm, limits treatment options when needed. Better *S*.Tm control would therefore be of benefit both for farm animals and for safety of the human food chain.

A promising pre-harvest intervention is vaccination. In this study we tested safety and immunogenicity of an oral inactivated *S*.Tm vaccine, which has been recently shown to generate an “evolutionary trap” and to massively reduce *S*.Tm colonization and transmission in mice. We show that this vaccine is highly immunogenic and safe in post-weaning pigs and that administration of a single oral dose results in a strong and long-lasting serum IgG response. This has several advantages over existing – mainly live – vaccines against *S*.Tm, both in improved seroconversion and reduced risk of vaccine-strain persistence and reversion to virulence.

## Introduction

The gram-negative pathogen *Salmonella enterica* can cause infections in pigs that can manifest either as clinical disease or as subclinical carriage. Clinical disease, which – depending on the serotype – manifests as fibrinonecrotic enterocolitis or septicemia, causes morbidity and death in animals [1]. *Salmonella* carriage, including symptomless colonization, is a risk for transmission both to other pigs and along the foodchain [2, 3]. Stressors such as transport and diet shifts or food deprivation increase both shedding and susceptibility of pigs to infection with *Salmonella* driving horizontal transmission between pigs at times when herd mixing is most likely. This corresponds with peaks of clinical disease on transfer between farrowing/fattening/finishing farms. This also increases the risk of carcass contamination after transport to the abattoir. Vertical transmission from mother to offspring is another common transmission route ensuring *Salmonella* persistence on a herd level [2]. In the EU, salmonellosis is the second most common zoonosis and 12% of the around 90 000 yearly cases of human salmonellosis originate from pork and pork products [4]. Almost all such cases are caused by *Salmonella enterica* subspecies *enterica* serovar Typhiumurium (*S*.Tm) [4]. Therefore, strategies to not only prevent clinical disease, but also to eliminate subclinical carriage of *Salmonella* in swine are urgently required.

Next to antibiotics, vaccines are one of the most successful interventions against bacteria [5]. Current *S*.Tm vaccinations have some efficacy in disease prevention and variable efficacy in reduction of shedding, but they do not decolonize the herd. Correspondingly, it remains impossible to completely exclude *Salmonella* from a farm with known carriage, without culling all animals [6-10]. In a model mimicking pre-slaughter stress, fecal shedding was not different between pigs vaccinated with the licensed Salmoporc vaccine and unvaccinated animals [3]. Moreover, attenuated live vaccines have been found to be themselves readily shed and can cause disease in permissive conditions such as reduced intestinal microbial diversity or in immunocompromised hosts [11, 12].

In mice, we have recently demonstrated that rationally-designed inactivated whole-cell oral vaccines “EvoTrap vaccines” can induce an intestinal antibody response, which directs evolution of loss-of-virulence of *Salmonella* [13]. In contrast, vaccination with an inactivated wild-type *S*.Tm (*S*.Tm^WT^) vaccine leads to the emergence of escaper mutants with equal fitness and equal virulence. EvoTrap vaccination achieves both a better disease outcome and greatly reduced transmission of *S*.Tm in a non-Typhoidal Salmonellosis mouse model [13]. The EvoTrap vaccine consists of four peracetic acid (PA) inactivated *S*.Tm strains comprising all *S*.Tm O-antigen variants (O:4[5],12-0, O:4,12-0, O:4[5],12-2, O:4,12-2). This induces an intestinal IgA response that prevents easy immune escape of the bacteria from the antibody response via O-antigen modification (e.g. changing the acetylation status of Abequose (O:4→O:5) [14] or by glucose addition to the backbone galactose residue (O:12-0→O:12-2) [15]). Instead, EvoTrap vaccination in mice forces an evolutionary trade-off in *Salmonella*, via antibody-mediated selection of mutants that have lost production of the polymerized O-antigen. These arising escaper mutants carry a major genomic deletion, mediated by flanking inverted repeats, that includes the O-antigen polymerase gene *wzyB* (also known as *rfbP*) [13]. O-antigen loss compromises membrane stability, making *S*.Tm^wzyB^ more susceptible to complement, bile acids, bacteriophage predation and different other environmental stressors and thereby dramatically reducing transmission [13]. As these inactivated oral vaccines are cheap and easy to apply, and are expected to have excellent safety profiles, we here set out to test whether such vaccines are safe and immunogenic in domestic pigs.

Here we show that oral vaccination of domestic pigs with an inactivated *S*.Tm^WT^ vaccine (O:4[5],12-0) and the EvoTrap vaccine is safe and induces a strong *S*.Tm-specific IgG response in serum. Vaccination with the EvoTrap vaccine results in antibodies directed against all known *S*.Tm O-antigen variants and a single oral dose is sufficient to induce this response. The *S*.Tm-specific IgG titre in pigs vaccinated just once is still 100-fold higher than in naïve pigs two months after vaccination.

## Methods

### Strains and plasmids

All strains used in this study are listed in the Table 1.

**Table 1.**
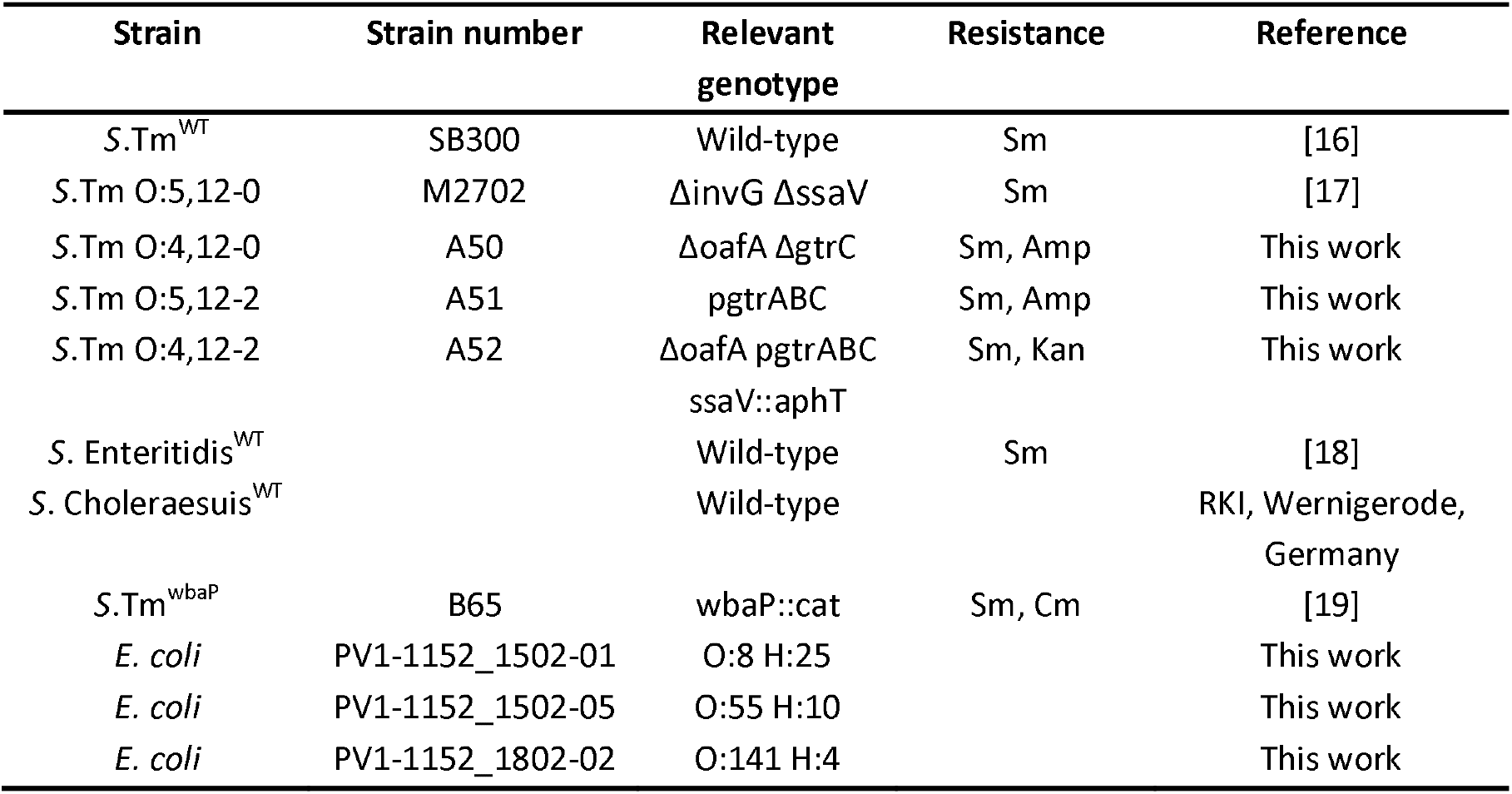
Strains used in this study.

**Table 2.** Composition of medium for high cell density cultivation of *S*.Tm.

| Component | Batch medium | Feeding solution |
| --- | --- | --- |
| D-(+)-Glucose | 27.72 g l<sup>-1</sup> | 770 g l<sup>-1</sup> |
| (NH<sub>4</sub>)<sub>2</sub>HPO<sub>4</sub> | 5.75 g l<sup>-1</sup> | - |
| KH<sub>2</sub>PO<sub>4</sub> | 7.69 g l<sup>-1</sup> | - |
| MgSO<sub>4</sub> · 7H<sub>2</sub>O | 1.23 g l<sup>-1</sup> | 19.96 g l<sup>-1</sup> |
| NH<sub>4</sub>Cl | 5.35 g l<sup>-1</sup> | 64.18 g l<sup>-1</sup> |
| L-Histidine | 1.55 g l<sup>-1</sup> | 6.2 g l<sup>-1</sup> |
| Fe(III)-citrate | 100 mg l<sup>-1</sup> | 0.16 g l<sup>-1</sup> |
| CoCl<sub>2</sub> | 2.5 mg l<sup>-1</sup> | 4 mg l<sup>-1</sup> |
| MnCl<sub>2</sub> | 15 mg l<sup>-1</sup> | 24 mg l<sup>-1</sup> |
| CuCl<sub>2</sub> | 1.5 mg l<sup>-1</sup> | 2.4 mg l<sup>-1</sup> |
| H<sub>3</sub>BO<sub>3</sub> | 3 mg l<sup>-1</sup> | 4.8 mg l<sup>-1</sup> |
| Na<sub>2</sub>MoO<sub>4</sub> · 2H<sub>2</sub>O | 2.5 mg l<sup>-1</sup> | 4 mg l<sup>-1</sup> |
| Zn(CH<sub>3</sub>COO)<sub>2</sub> · 2H<sub>2</sub>O | 13 mg l<sup>-1</sup> | 20.8 mg l<sup>-1</sup> |
| EDTA | 8.4 mg l<sup>-1</sup> | 13.44 mg l<sup>-1</sup> |
| H<sub>2</sub>SO<sub>4</sub> | - | 0.5 M |
| NH<sub>4</sub>OH | - | 4 M |
| Antifoam-C | - | 1% (v/v) |

All *Salmonella* and *E. coli* were cultivated in lysogeny broth (LB) or high cell density cultivation medium containing appropriate antibiotics (100 µg/ml streptomycin (AppliChem); 15 µg/ml chloramphenicol (AppliChem); 50 µg/ml kanamycin (AppliChem); 50 µg/ml ampicillin (AppliChem)). Dilutions were prepared in Phosphate Buffered Saline (PBS, Difco).

### Media

High cell density cultivation medium was adapted from Riesenberg *et al*. [20] to be suited for *S*.Tm cultivation.-

Medium was prepared as previously described [20].

### Vaccine production

Bacteria for small batch vaccine production were grown in LB containing the appropriate antibiotics. If more vaccine was required, bacteria grown in LB to late logarithmic phase were subcultured overnight in high cell density cultivation medium before being transferred to a 3.6 l bioreactor (Infors) for fed-batch cultivation. Inoculation of the bioreactor was typically done with an OD_600_ of 0.1 into the initial culture volume of 1.5 l. Main cultivations were done at 37 °C with a pH set-point of 7.4. Stirring speed and air flow were adjusted to keep the partial oxygen pressure at around 20%. Upon rapid oxygen increase – marking bacterial growth arrest due to nutrient depletion – feeding solution was added manually until the desired biomass was reached. Cultures were harvested at an OD_600_ of 25-35 and used for vaccine production. The bioreactor was refilled with fresh high cell density cultivation medium until the final number of bacterial cells was reached. For overnight growth, the temperature was reduced to 25 °C, the air flow to 1.5 l min^-1^ and the stirrer speed to 800 rpm.

Harvested bacteria were killed with peracetic acid (PA) as previously described [21]. Briefly, bacteria were harvested by centrifugation and re-suspended to a density of 10^9^-10^10^ per ml in sterile PBS or directly from the bioreactor. Peracetic acid (Sigma-Aldrich) was added to a final concentration of 0.4% v/v. The suspension was mixed thoroughly and incubated for 60 min at room temperature. Bacteria were washed with sterile PBS until the resulting vaccine had a pH between 6 and 7. The final pellet was resuspended to yield a density of 10^11^-10^12^ particles per ml in sterile PBS. The exact number was determined by flow cytometry using counting beads (Fluoresbrite® Multifluorescent Microspheres). Liquid vaccines were stored at 4 °C for up to three weeks or lyophilized and stored at room temperature. Each batch of vaccine was tested for sterility and correct composition of O-antigens before use.

### Pig vaccinations

All animal experiments were performed in accordance with Swiss Federal regulations approved by the Commission for Animal Experimentation of the Kanton Zurich (licenses 077/2018 and 016/2021; Kantonales Veterinäramt Zürich, Switzerland). Conventionally reared, outbred piglets with different genetic contributions of French-Swiss Landrace, Swiss Large White and Duroc were used. All piglets were either bred at Strickhof, Lindau, Switzerland or at the Department for Farm Animals, Vetsuisse Faculty, Zurich, Switzerland. Piglets were weaned at 3-4 weeks and transported to the experimental facility. Piglets were fed ad libitum with a comercial diet (Granovit® Ferkelfutter Classic). Vaccinations started at an age of 4-5 weeks after one week of acclimatization and training. Vaccinations were done by administering 5∙10^8^ bacterial particles intranasally or 5∙10^11^ or 8∙10^12^ orally as stated in the text and figures. Where multiple strains were used, equal numbers of each strain were given. All animals included in experiments were objectively healthy as determined by routine health checks. Piglets were randomly assigned to experimental groups and wherever possible an equal number of castrated males and females was used in each group. Researchers were not blinded to group allocation because most readouts were quantitative and not subjective.

### Analysis of specific antibody titres by bacterial flow cytometry

Specific antibody titres in serum were measured by flow cytometry as described [22]. Briefly, blood was collected from the jugular vein into a Sarstedt S-Monovette® Serum 9 ml tube. Blood was centrifuged at 2000x *g* for 10 min to obtain serum, heat-inactivated at 56 °C for 30 min and stored at -20 °C until further analysis. Bacterial targets (antigen against which antibodies are to be titred) were grown overnight in LB, then gently pelleted for 2 min at 7000x *g*. The pellet was washed with 0.2 µm-filtered PBS before resuspending at a density of approximately 10^7^ bacteria per ml. Serum was thawed and used to perform serial dilutions starting from 1:10. 50 μl of the dilutions were incubated with 50 μl bacterial suspension for 15 min at room temperature. Bacteria were washed twice with 150 μl PBS by centrifugation at 7000x *g* for 5 min, before resuspending in 25 μl of 0.2 µm-filtered PBS containing polyclonal biotinylated Anti-Pig IgG (Serotec, 5 µg/ml, AB_10673135). After 30 min of incubation at 4 °C, bacteria were washed twice with PBS as above and stained with Streptavidin-APC (BioLegend, 3.3 µg/ml) for 30 min at 4 °C. After washing twice, bacteria were resuspended in 100 μl PBS for acquisition on a Beckman Coulter Cytoflex S. Data were analysed using FlowJo (Treestar). After gating on bacterial particles, log-median fluorescence intensities (MFI) were plotted against serum dilution factor for each sample and 4-parameter logistic curves were fitted using Prism with the bottom fixed to the average MFI of control samples without serum (Graphpad, USA). Titers were calculated from these curves as the dilution factor giving an above-background signal.

### Cross-absorption

Bacteria were grown overnight in LB, washed twice with PBS and resuspended at 10^9^ bacteria per ml. Heat-inactivated serum samples were diluted 1:5 (i.e. double of the starting dilution used for flow cytometry) and incubated with an equal volume of bacteria for 1 h at 4 °C with gentle shaking. Bacteria were removed by centrifugation for 5 minutes at 17000x *g* and filtration through a 0.22 μm filter. Cross-absorbed serum samples were then used for flow cytometry as described above.

### Histological procedures

Tissue embedded in paraffin was cut into 2 μm cryosections and mounted on glass slides. Cryosections were air dried overnight and stained with hematoxylin and eosin (H&E).

### Analysis of O-antigen composition by bacterial flow cytometry

Single EvoTrap vaccine strains or the corresponding *S*.Tm overnight cultures were diluted to app. 10^7^ cells per ml in 0.2 µm-filtered filtered PBS. Staining was performed with 0.2 µm-filtered solutions of STA5 (human recombinant monoclonal IgG2 anti-O:12-0, 3.2 µg/ml) (Moor et al., 2017) or rabbit anti-Salmonella O:5 (Difco, 1:200, 226601) for 30 min at 4 °C. Bacteria were washed twice with filtered PBS and then resuspended in 0.2 µm-filtered solutions of appropriate secondary reagents (Alexa 647-anti-human IgG (Jackson ImmunoResearch, 1:100, 109-605-098, AB_2337889) and Brilliant Violet 421-anti-rabbit IgG (BioLegend, 1:100, 406410, AB_10897810)). This was incubated for 30 min at 4 °C before the cells were washed as above and resuspended for acquisition on a Beckman Coulter Cytoflex S. Data were analysed using FlowJo (Treestar).

### Salmonella enrichment cultures

Pig feces was regularly sampled from individual pigs and from the pen and tested for the presence of *Salmonella*. Feces was pre-cultured in phosphate buffered water to enrich for Enterobacteriaceae and 100 μl were transferred into selenite cysteine broth for specific *Salmonella* enrichment. Cultures were then plated on MacConkey agar and non-lactose fermenting single colonies send for 16S sequencing.

### Statistical analysis

Researchers were not blinded for the assignment of the experimental groups and the data analysis. Kruskal-Wallis test followed by Dunn ‘s post hoc test was used for comparison of three groups. Statistical analysis was performed with Graphpad Prism Version 9.2.0 for Windows (GraphPad Software, La Jolla, California USA). All p values were reported.

## Results

### An oral inactivated bacterial vaccine shows excellent safety and efficacy in pigs

While inactivated oral vaccines have shown excellent immunogenicity and safety in murine models, porcine biology is considerably different [23]. We therefore first designed experiments to test the optimal dosing of inactivated oral vaccines in 4–12-week-old pigs, with a primary readout of weight gain and *Salmonella-*specific antibody titres.

Our main goal was to create a new pig vaccine with increased safety and efficacy in comparison to already existing vaccines. Therefore, a major focus of our initial experiments was vaccine safety. For this reason, all piglets were monitored daily for any adverse effects and for weight gain during the vaccination period with the inactivated PA-*S*.Tm^WT^ vaccine. In addition, two pigs receiving the standard dose of 5∙10^11^ PA-*S*.Tm^WT^ were monitored for almost two more months after the completion of the normal vaccination scheme of six doses in weekly intervals. All animals were in good health and showed robust weight gain during and after PA-*S*.Tm^WT^ vaccination independent of vaccine dosage (**Fig. 1A**). Occasionally, piglets had slight diarrhea, which was however not more frequent than in untreated post-weaning piglets. We did not expect to see any reduction in the frequency of diarrhea due to vaccination as our experiments were performed on a *Salmonella*-free farm and on piglets born to *S*.Tm antibody negative sows (data not shown) i.e. *Salmonella* was not expected to be a causative agent of diarrhea in this herd. To further corroborate our findings of no adverse effect, we performed a comprehensive histological evaluation on the two pigs that were observed for longer. To also exclude more acute effects of our vaccine, both pigs received a further vaccine dose five days before euthanasia. Histopathological evaluation of heart, lung, liver, spleen, kidney and intestine (duodenum and colon) revealed good health of all animals (**Fig. 1B**). To evaluate the immunogenicity of our novel *Salmonella* vaccine, we checked the serum of vaccinated pigs for *S*.Tm-specific IgG by flow cytometry [22]. All vaccinated piglets had no or background response to *S*.tm before vaccination (**Fig. 1C**). Vaccination with both the standard (5∙10^11^) and the high dose (8∙10^12^) induced a strong *S*.Tm specific serum IgG response, which was absent in untreated age-matched piglets from the same swine herd (**Fig. 1C, D**). We did not see any difference in antibody titres between the lower and the higher dose groups, indicating that we had already saturated the system with 10^11^ vaccine particles. To further check that the observed IgG response is indeed vaccine-induced and specific for *S*.Tm, we performed cross-absorption of the pig serum against *Salmonella* Enteritidis (O:1,9,12), *Salmonella* Choleraesuis (O:6,7), an O-antigen deficient mutant *S*.Tm^wbaP^, as well as a mix of three *E. coli* strains (O:8 H:25, O:55 H:10, O:141 H:4) isolated from the same pig farm. None of these treatments reduced the detectable antibody response against *S*.Tm^WT^ in the vaccinated pigs, while cross-absorption against the vaccine-strain *S*.Tm^WT^ (O:5,12-0) reduced antibody-binding to naïve levels (**Fig. S1**). We therefore conclude that the induced antibodies are highly specific to the *S*.Tm O-antigen present in our vaccine.

**Figure 1.**
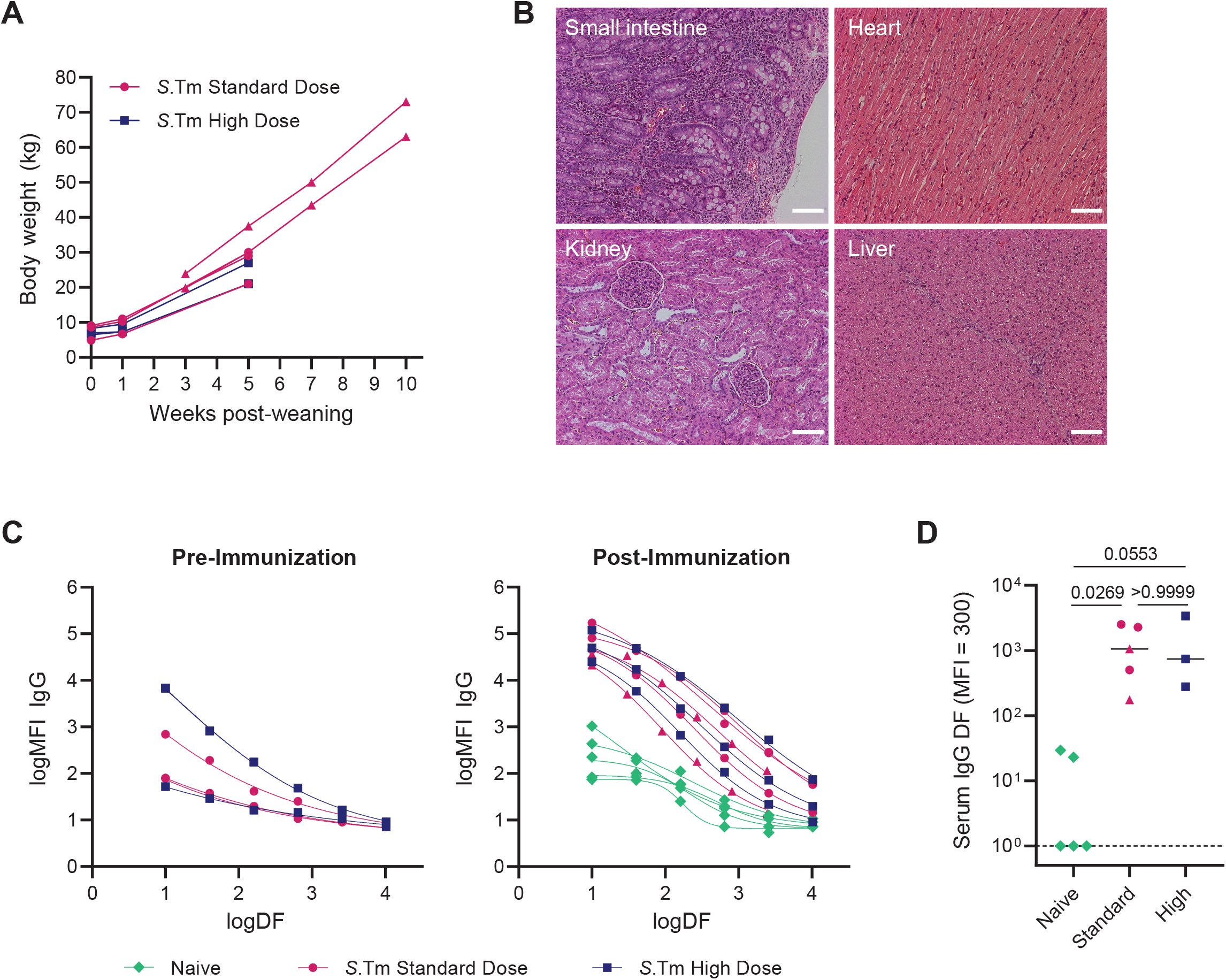
Oral vaccination with peracetic acid-inactivated *S*.Tm (PA-*S*.Tm^WT^) demonstrates excellent safety and efficacy in pigs. Pigs were orally vaccinated 6 times in weekly intervals starting at an age of 4 weeks. Pigs received either 5∙10^11^ (*S*.Tm Standard Dose, pink symbols) or 8∙10^12^ (*S*.Tm High Dose, blue squares) PA-*S*.Tm^WT^. (**A**) All vaccinated pigs showed normal weight gain during the vaccination period and afterwards. (**B**) Representative histopathology of H&E-stained sections from small intestine, heart, kidney and liver revealed robust health in vaccinated animals. Scale bars: 100 µm. (**C**) **S**.Tm-specific serum IgG was determined by flow cytometry prior to vaccination and one week after completion of the whole vaccination scheme. The post-immunization response was furthermore compared to naïve age matched pigs from the same herd (Naive, green diamonds). (**D**) Post-immunization titres calculated from the curves in (C). Triangles show pigs that were followed for 8 more weeks after the last standard vaccination. These pigs received an additional dose of vaccine to see also possible acute effects in histology. Antibody titres were determined one week after the standard vaccination scheme (6 doses) for all pigs. Solid lines depict the median, dotted lines show the detection limit. Statistics were calculated with Kruskal-Wallis test. DF, dilution factor; MFI, median fluorescence intensity.

Taken together, we could show that oral vaccination with 10^11^ particles of our inactivated oral vaccine has an excellent safety profile and induces a strong and highly specific serum IgG response. Of note, we tried extensively to measure specific IgA responses in pig intestinal samples. However, in the absence of any positive control we cannot currently differentiate between poor performance of the anti-pig-IgA antibodies or actual absence of an intestinal IgA response.

### EvoTrap vaccination leads to IgG induction specific for all possible.Tm O-antigens

As our first experiments confirmed good safety and induction of a highly specific IgG response against the *S*.Tm O-antigen, we expanded these observations to the complete EvoTrap oral vaccine. Intranasal vaccination typically requires lower doses than oral vaccination, which would be beneficial in reducing vaccine costs and would potentially allow administration to big numbers of animals via aerosol inhalation. Previous work in mice demonstrated effective cross-priming between intranasal and intestinal sites [24]. We therefore tested the potential of a 1000-fold lower vaccine dose administered via the intranasal route to boost oral vaccination (**Fig. 2A**).

**Figure 2.**
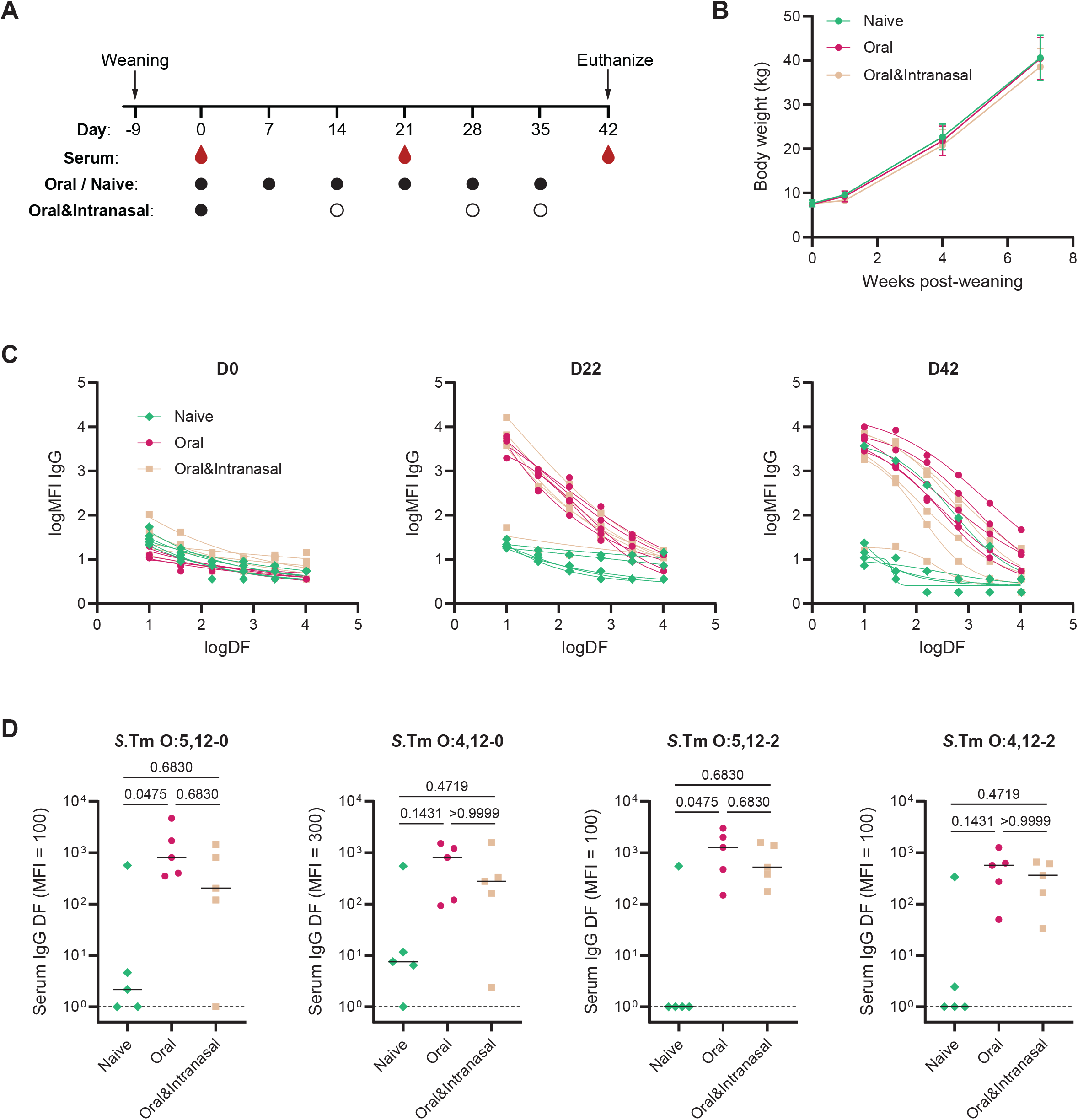
EvoTrap vaccination leads to IgG induction specific for all possible *S*.Tm O-antigens. Pigs were either orally vaccinated for 6 times in weekly intervals with the EvoTrap vaccine (5∙10^11^ per strain, pink circles) or mock-vaccinated with vaccine carries (green diamonds). A third group was orally primed with the EvoTrap vaccine (5∙10^11^ per strain), followed by three intranasal boosters (5∙10^8^ per strain, beige squares). (**A**) Experimental procedure. Black circles indicate oral vaccinations, white circles show intranasal vaccinations. (**B**) Weight gain was equal for all groups. Mean±SD. (**C**) **S**.Tm O:5,12-0-specific serum IgG was determined by flow cytometry prior to vaccination, midway and one week after completion of the whole vaccination scheme. (**D**) Post-immunization titres against all *S*.Tm O-antigens. Solid lines depict the median, dotted lines show the detection limit. Statistics were calculated with Kruskal-Wallis test. DF, dilution factor; MFI, median fluorescence intensity.

Again, all three groups showed an equal weight gain and no vaccine associated adverse effects (**Fig. 2B**). Blood sampling was done on the day of the first vaccination (D0), one week after the third and second vaccination, respectively, (D22) and one week after completion of the full vaccination scheme (D42, **Fig. 2A**). Both vaccinated groups showed the induction of an O:5,12-0-specific IgG response at D22. This antibody response was found to be further enhanced on completion of the full vaccination scheme (**Fig. 2C**). Interestingly, one pig from the control group showed a strong O:5,12-0-specific IgG response at D42, which was completely absent on D22. The reasons for this are unclear. Our detection limit for subclinical infection with *S*.Tm was as low as 10^3^ bacteria/ml feces and no sample ever tested positive. Nevertheless, we cannot exclude very low-level colonization or very transient infection between samplings, or cross-reactivity to a different invasive microbiota member. When checking endpoint responses to all four possible *S*.Tm O-antigens, we found them to be greatly enhanced in both vaccination groups, with one piglet from the oral-intranasal group showing a highly biased response against O:12-2-carrying *S*.Tm strains. (**Fig. 2D**).

In summary, we could show the induction of a robust IgG response against all possible *S*.Tm O-antigens by using the EvoTrap vaccine. Moreover, intranasal boosting seems to be a good possibility to decrease the required amount of vaccine.

### A single oral EvoTrap dose is sufficient for the induction of a strong and long-lived antibody response

Weekly vaccine feeding was initially chosen to mimic the most common protocol used in mice. However, this dosing is not necessarily practical in the context of a working farm, and the cost scales with the number of boosters administered. To further reduce the amount of vaccine needed per animal, we tested whether a single EvoTrap vaccination or vaccination followed by one booster would be sufficient to induce a good antibody response. Therefore, piglets were vaccinated either 1) once orally with 5∙10^11^ bacteria per strain, 2) once intranasally with 5∙10^8^ bacteria per strain, or 3) primed intranasally with 5∙10^8^ bacteria per strain followed by an oral booster vaccination with 5∙10^11^ bacteria per strain two weeks later (**Fig. 3A**). We observed that a single oral EvoTrap dose was sufficient to induce a *S*.Tm-specific serum IgG, while this was not the case for only intranasal vaccination. Intranasal priming in combination with an oral booster led to a similar response as the oral vaccination alone (**Fig. 3B**). We further checked the longevity of the induced serum IgG response over two months following the first vaccination. The initially high antibody levels waned slowly over time but were still easily detectable in all orally-vaccinated animals 2 months after the first vaccination (**Fig. 3B**). This was true for all *S*.Tm O-antigens tested (**Fig. 3B**).

**Figure 3.**
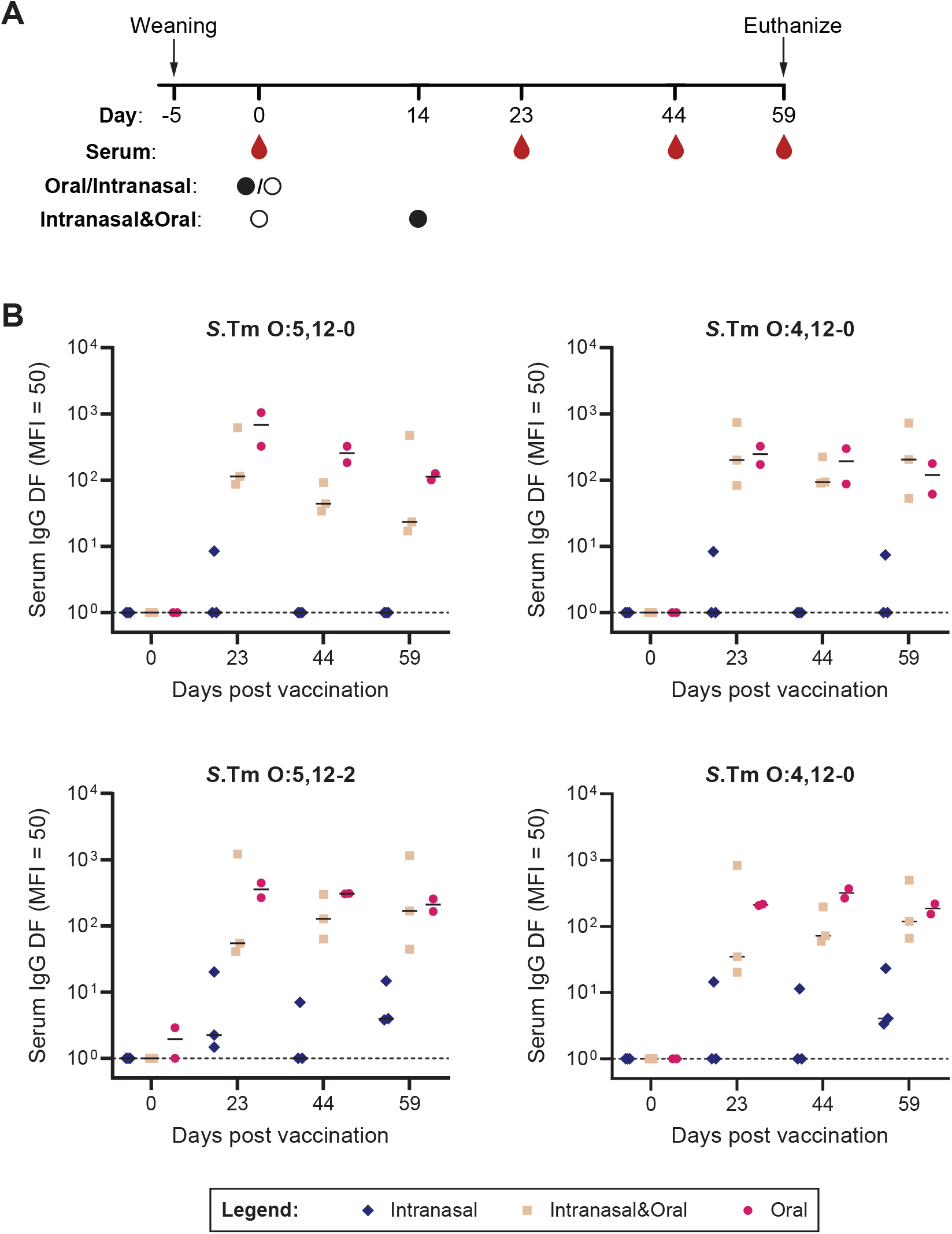
A single oral EvoTrap dose is sufficient for the induction of a strong and long-lived antibody response. Pigs were either orally (5∙10^11^ per strain, pink symbols) or intranasally (5∙10^8^ per strain, blue symbols) vaccinated with EvoTrap vaccine or intranasally primed followed by an oral booster two weeks later (beige symbols). (**A**) Experimental procedure. Black circles indicate oral vaccinations, white circles show intranasal vaccinations. (**B**) Serum IgG titres over time against all *S*.Tm O-antigens as determined by flow cytometry. Solid lines depict the median, dotted lines show the detection limit. DF, dilution factor; MFI, median fluorescence intensity.

Taken together, we could show that a single oral dose is sufficient for the induction of a strong *S*.Tm-specific IgG response, while this cannot be achieved with intranasal vaccination alone. This antibody-response is long-lived.

## Discussion

Inactivated bacterial vaccines have often been considered poorly immunogenic [25]. We could show that this is not the case for both of our orally administered inactivated *S*.Tm vaccines in two different cohorts of domestic pigs in Switzerland. Even a single dose of EvoTrap vaccine given without any additional adjuvants could induce a strong, highly specific and long-lasting serum IgG response against *S*.Tm. In contrast, the only licensed *S*.Tm vaccine in Europe, which is a live-attenuated vaccine, failed to induce IgG above the level of control animals within 3 weeks after a single oral vaccination [26]. Combined with potential safety risks of live-vaccines, including prolonged carriage and shedding, this suggests the potential for inactivated oral vaccines in improving swine health. The inactivated vaccine also showed good stability upon lyophilization and storage at room temperature, and is therefore practical to distribute, store and feed (**Fig. S2**). By this we also circumvent another problem of live vaccines, namely degradation or inactivation by inappropriate storage or interfering substances such as feed additives, which could accidentally kill the vaccine strain.

Although robust serum antibody responses could be measured with our vaccine, it remains unclear whether this can translate into effective protection. Several studies using live-attenuated or inactivated vaccines have demonstrated reduction of Salmonella or even complete protection from environmental *S*.Tm infection [27-30]. Moreover, the importance of involvement of a mucosal immune response was underscored by the fact that reduction in the percentage of shedder sows and subsequent complete protection of piglets from acquiring *S*.Tm was seen only in the study that used oral priming (in combination with i.m. boosters) on a farm with endemic vaccine-identical strain [27]. Autogenous vaccination requires individual diagnosis and vaccine development, which is time consuming and costly. Therefore, vaccination regimens that induce a broader protection would be preferable. Time and route of vaccination can have an impact on protective efficacy. Although different routes and intervention regimens (sows, newborn piglets, weaners, fattening pigs) and combinations thereof have been tested in the past, it is difficult to compare these studies as they differ in a variety of other factors (e.g. vaccine used, mode of infection, readout) [8]. Although further work, and challenge studies, are still needed to determine the best timing, dose and route of our novel *Salmonella* vaccine to achieve optimal protection, we have proofed safety and immunogenicity. In this study we did not measure the induction of cell mediated immunity, who ‘s induction typically observed to be stronger with live vaccines. Cell-mediated immunity was also found to confer cross-protection [31] and should be checked in subsequent studies that also quantify protection.

Based on the current study design a conclusion if the immune response was induced by the intranasal application or by the oral booster is not possible and requires further investigation (**Fig. 3**). Blood sampling between prime and boost vaccination was not possible for the required vaccination scheme due to animal license restrictions. However, we could see an antibody increase from D22 to D42 in our oral-primed and intranasally boosted group (**Fig. 2**), hinting at a small effect of intranasal vaccination. It remains possible that the intranasal dose chosen was too low, and this warrants further investigation.

Efforts have been ongoing to produce so called DIVA (differentiating infected from vaccinated animals) vaccines to allow testing for infection by serology. This has not been implemented in our strategy but can be achieved by deletion of a single surface protein e.g. ompD [7] that is immunogenic but not part of the protective immune response. However, it is also important to note that serology does not give any information about the current state of infection and seropositive animals might not carry any *Salmonella* at the time point of testing. In fact, presence of *Salmonella*-specific antibodies has been found to poorly correlate with presence of bacteria in individual pigs [32].

Another demand for newly developed *Salmonella* vaccines is cross-protection. We have not tested this formally, but cross-absorption showed that the vaccination with PA-*S*.Tm induced highly *S*.Tm^WT^ O-antigen specific antibodies. However, the EvoTrap vaccine induced antibodies against all possible full-length O-antigen variants found in *S*.Tm, which is the major serovar of concern for human infection in pig farming. Nevertheless, *S*. Choleraesuis causes more severe disease in pigs and O:6,7-producing strains should be included in a next generation of the vaccine in order to generate broader coverage.

Overall, we could demonstrate that EvoTrap vaccination is highly efficient in generating a serum IgG response upon oral administration to pigs. A single oral dose of the EvoTrap vaccine was sufficient to induce a high titre of *S*.Tm-specific IgG, which has not been possible with the currently licensed porcine *S*.Tm vaccine in a previous study [26].

## Supporting information

Highlights

## Acknowledgements

We thank Salomé LeibundGut-Landmann for giving us unrestricted access to laboratory facilities at the VetSuisse Faculty. This has immensely facilitated immediate sample processing. We thank Mick Bailey for sharing his knowledge about pig mucosal immunology and behaviour, giving us a head start in our first practical experiments. We thank Nadja Aeberhard, Patricia Sutter and Ramona Wissmann for their veterinary support at blood sampling and during dissection. We also want to thank the staff at the Department for Farm Animals, Vetsuisse Faculty for their excellent support. We thank Frauke Seehusen from the LAMP for assistance with histopathology.

Funding for this work was provided by the Gebert Rüf Microbials (GR073_17). VL, and ES are supported by the Gebert Rüf Microbials (GR073_17). ES acknowledges the support of the Swiss National Science Foundation (40B2-0_180953, 310030_185128), and European Research Council Consolidator Grant (865730). This work was supported as a part of NCCR Microbiomes, a National Centre of Competence in Research, funded by the Swiss National Science Foundation (grant number 180575). Funding was provided by the Botnar Research Centre for Child Health as part of the Multi-Investigator Project: Microbiota Engineering for Child Health.

## Author contributions

Conceptualization, V.L., E.S.; Investigation, V.L., S.A. T.E., S.P., E.C.B., D.H., C.M.; Methodology, V.L.; Validation, V.L., S.A.; Data Curation, V.L., S.A..; Formal Analysis, V.L.; Project administration, V.L., E.S.; Funding acquisition E.S.; Resources, T.E., D.K., E.S.; Supervision, V.L., E.S.; Visualization, V.L., S.A..; Writing – original draft, V.L., E.S.; Writing – review and editing, V.L., S.A., T.E., S.P., C.M., D.K., E.S.;

## Supplementary figure titles and legends

**Figure S1.**
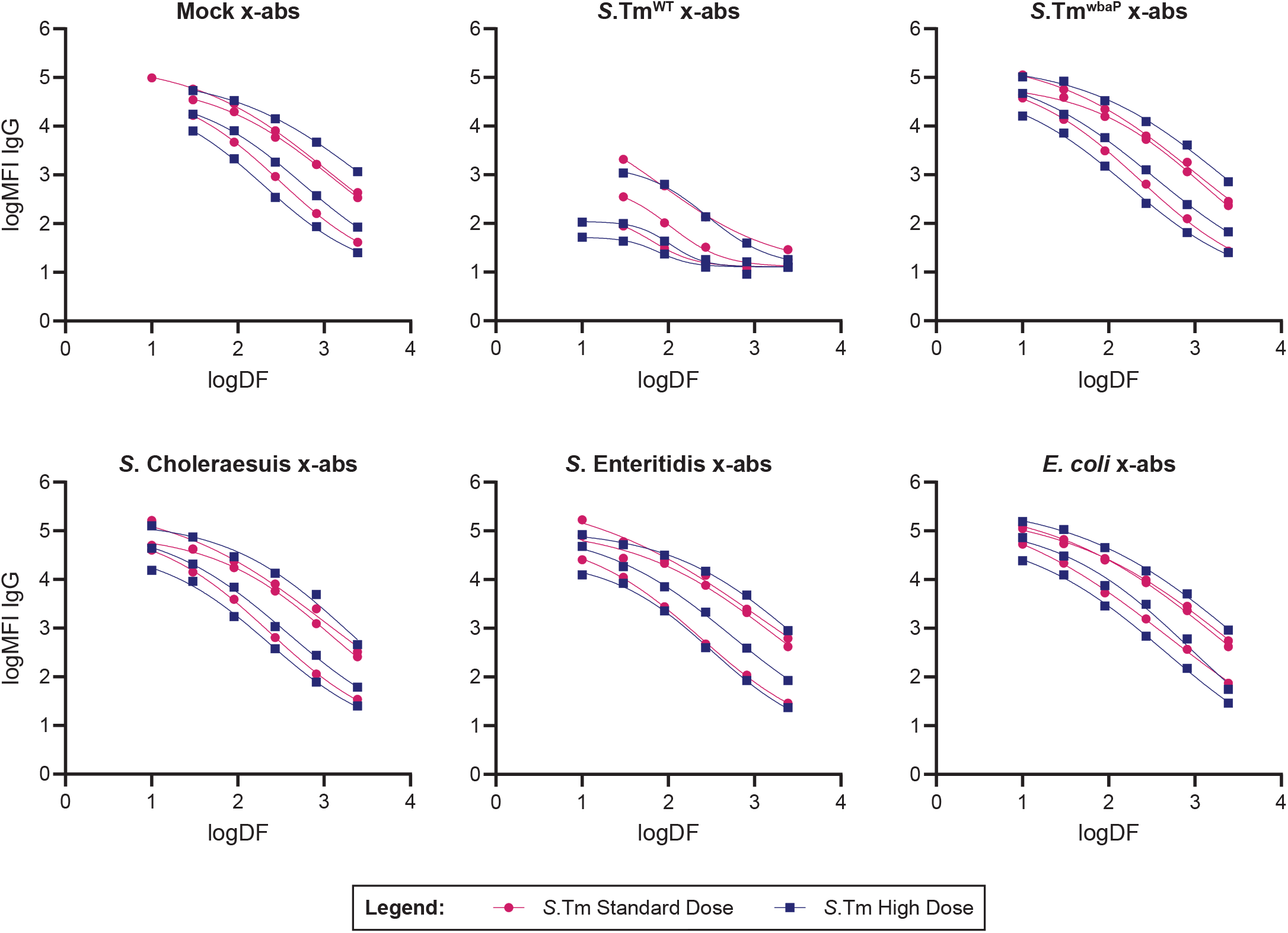
The induced antibody response is highly specific for the *S*.Tm O-antigen (related to Fig. 1). Pigs were orally vaccinated 6 times in weekly intervals starting at an age of 4 weeks. Pigs received either 5∙10^11^ (*S*.Tm Standard Dose, pink circles) or 8∙10^12^ (*S*.Tm High Dose, blue squares) PA-*S*.Tm. Specificity of serum IgG was determined by cross-absorption with different Salmonella serovars (*S*.Tm^WT^, *S*. Choleraesuis, *S*. Enteritidis), a mix of three *E. coli* strains (O:8 H:25, O:55 H:10, O:141 H:4) isolated from the pig herd or the O-antigen deficient mutant *S*.Tm^wbaP^. DF, dilution factor; MFI, median fluorescence intensity.

**Figure S2.**
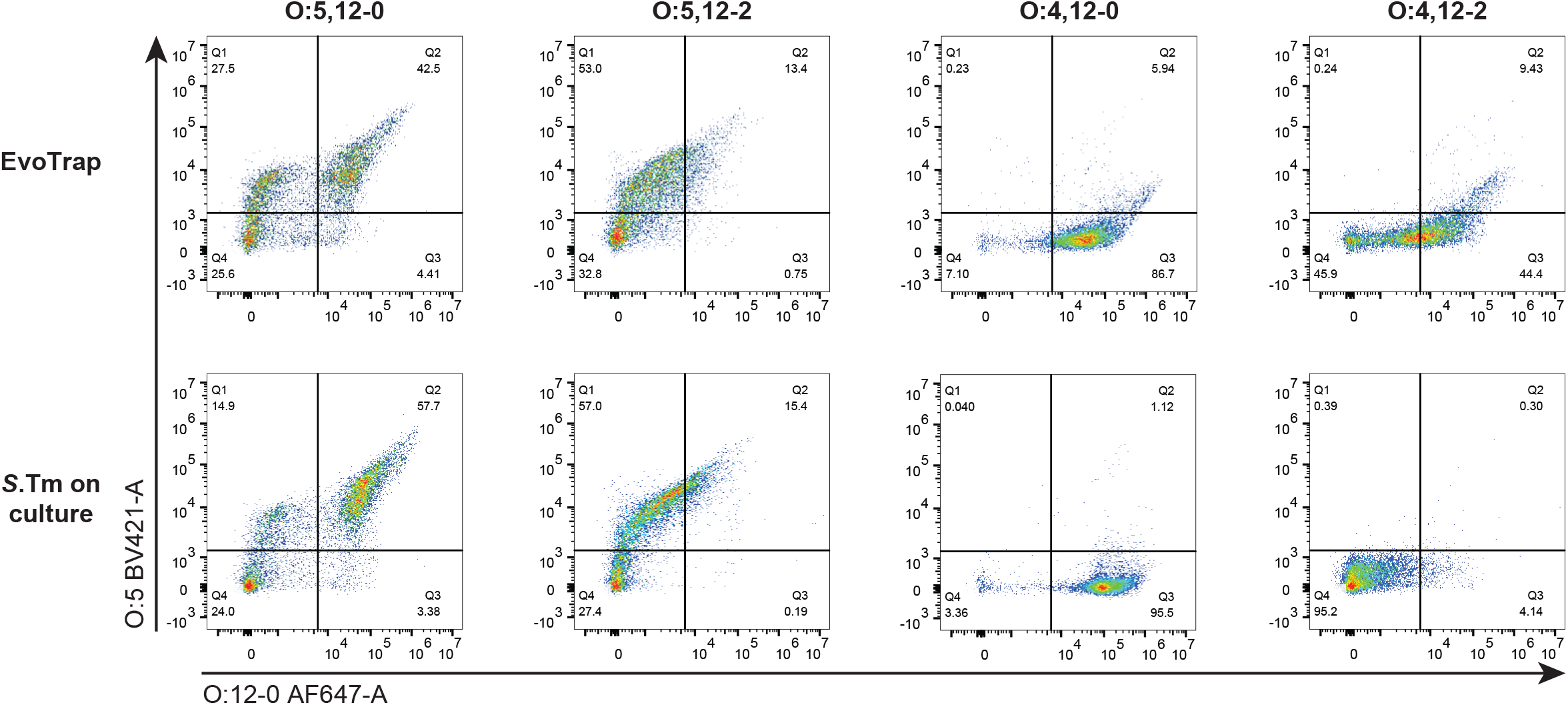
The EvoTrap vaccine shows good stability upon lyophilization and storage at room temperature for over a year. EvoTrap vaccine stability was analyzed by O:5 and O:12-0 staining in flow cytometry for single EvoTrap strains (upper panels) and corresponding fresh cultures of the respective *S*.Tm strains (lower panels).

