## Supplementary material for "A rationally designed inactivated *Salmonella* Typhimurium vaccine induces strong and long-lasting immune responses in pigs": Highlights

- An oral inactivated *Salmonella* Typhimurium vaccine shows excellent safety in pigs
- Vaccination induces *Salmonella* Typhimurium (*S*.Tm) O-antigen specific serum IgG
- A single dose is sufficient to induce a strong and long-lasting IgG response
- All *S*.Tm O-antigen variants are targeted by vaccination with the EvoTrap vaccine
